# Studying 3D genome evolution using genomic sequence

**DOI:** 10.1101/646851

**Authors:** Raphaël Mourad

**Affiliations:** Laboratoire de Biologie Cellulaire et Moléculaire du Contrôle de la Prolifération (LBCMCP), CNRS, Université Paul Sabatier (UPS), 31000 Toulouse, France

## Abstract

The 3D genome is essential to numerous key processes such as the regulation of gene expression and the replication-timing program. In vertebrates, chromatin looping is often mediated by CTCF, and marked by CTCF motif pairs in convergent orientation. Comparative Hi-C recently revealed that chromatin looping evolves across species. However, Hi-C experiments are complex and costly, which currently limits their use for evolutionary studies over a large number of species. Here, we propose a novel approach to study the 3D genome evolution in vertebrates using the genomic sequence only, *e.g*. without the need for Hi-C data. The approach is simple and relies on comparing the distances between convergent and divergent CTCF motifs (ratio *R*). We show that *R* is a powerful statistic to detect CTCF looping encoded in the human genome sequence, thus reflecting strong evolutionary constraints encoded in DNA and associated with the 3D genome. When comparing vertebrate genomes, our results reveal that *R* which underlies CTCF looping and TAD organization evolves over time and suggest that ancestral character reconstruction can be used to infer *R* in ancestral genomes.

## 1 Introduction

Chromosomes are tightly packed in three dimensions (3D) such that a 2-meter long human genome can fit into a nucleus of approximately 10 microns in diameter [8]. Over the past years, the 3D chromosome structure has been comprehensively explored by chromosome conformation capture combined with high-throughput sequencing technique (Hi-C) at an unprecedented resolution [4,13,21]. Multiple hierarchical levels of genome organization have been uncovered. Among them, topologically associating domains (TADs) [4, 21] and chromatin loops [19] represent a pervasive structural feature of the genome organization. Moreover, functional studies revealed that spatial organization of chromosomes is essential to numerous key processes such as for the regulation of gene expression by distal enhancers [13] or for the replication-timing program [18].

A growing body of evidence supports the role of insulator binding proteins (IBPs) such as CTCF, and cofactors like cohesin, as mediators of long-range chromatin contacts [17,21,23]. In mammals, depletions of CTCF and cohesin decreased chromatin contacts [25]. Moreover, high-resolution Hi-C mapping has recently revealed that loops that demarcate domains were often marked by asymmetric CTCF motifs where cohesin is recruited [19]. These results support the extrusion loop model where CTCF and cohesin act together to extrude unknotted loops during interphase [20].

CTCF is an 11-zinc-finger (ZF) protein that is functionally conserved in vertebrates and *Drosophila* melanogaster [9, 11]. CTCF-binding sites and Hox gene clusters were shown to be closely correlated throughout the animal kingdom suggesting the conservation of the Hox-CTCF link across the Bilateria, as principal organizer of bilaterian body plans [9]. Comparative Hi-C further showed that CTCF motif position and orientation are conserved across species and that divergence of CTCF binding is correlated with divergence of internal domain structure [24]. These observations suggest that the genome could undergo a continous flux of local conformation changes by CTCF motif turnover that allow or prevent de novo enhancer-promoter interactions and misexpression [7]. Thus, the comparative analysis of CTCF-mediated looping across species is crucial to understand how gene expression or other key processes evolve. However, 3D genome analysis relies on complex and costly Hi-C experiments, which currently limits their use for evolutionary studies over a large number of species.

Here, we propose a novel approach to study the 3D genome evolution in vertebrates using the genome sequence only, *e.g*. without the need for Hi-C data. Therefore, this approach allows a comprehensive analysis of vertebrate 3D genomes whose number is exponentially increasing due to ungoing large sequencing projects such as the Vertebrate Genomes Project (VGP). The approach is simple and relies on comparing the distances between convergent and divergent CTCF motifs (ratio *R*). We show that *R* is a powerful statistic to detect CTCF looping encoded in the human genome sequence, thus reflecting strong evolutionary constraints encoded in DNA and associated with the 3D genome organization. Moreover, we found that *R* varies depending on the chromosome region, such as 3D (sub-)compartments, suggesting that *R* is not homogenous along the genome and might functionally define 3D chromatin state. When comparing *R* across vertebrates, our results reveal that the distance between convergent motifs which underly CTCF looping and TAD organization evolves over time and suggest that ancestral character reconstruction can be used to infer *R* in ancestral genomes.

## 2 Results and Discussion

### 2.1 CTCF-mediated looping in 3D and 1D genome point of view

In vertebrates, the 3D genome is organized in chromatin loops often mediated by CTCF and cohesin: the CTCF-mediated loops. In particular, CTCF sites at loop anchors occur predominantly (> 90%) in a convergent orientation, *i.e*. with a forward motif on the left anchor and a reverse motif on the right anchor [19] (Figure 1A). From a 1D genome point of view, the CTCF-mediated looping implies that two motifs in convergent orientation should be located farther apart than two motifs in divergent orientation (Figure 1B). Thus, based on this implication, we sought to compare the distances between contigous motifs depending on their orientation as a mean to study 3D genome from genomic sequence in species for which Hi-C data were not available.

**Figure 1.**
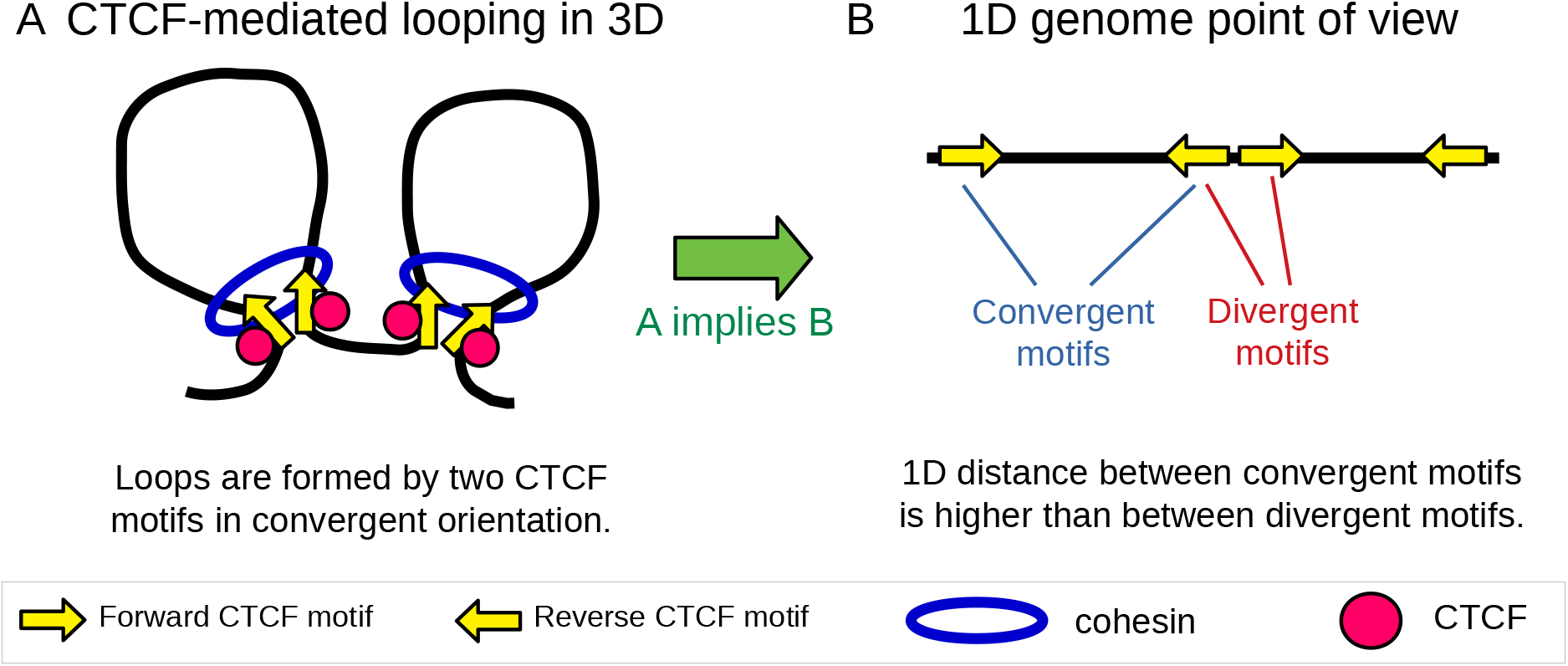
CTCF-mediated looping in 3D and 1D genome points of view. A) The CTCF-mediated looping in 3D. B) The 1D genome point of view of CTCF-mediated looping.

For this purpose, we estimated the following ratio *R*:

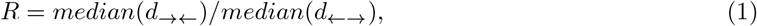

that was the ratio of two medians: the median of the distances between two contigous motifs in convergent orientation (noted “→ ←”), and the median of the distances between two contigous motifs in divergent orientation (noted “← →”). We hypothesized that the higher the ratio *R*, the higher CTCF looping in the genome. Because *R* was a ratio of distance medians, it accounted for the genome size effect and could thus allow comparisons between different genomes whose sizes may vary.

Additionally, we estimated another ratio used as a control:

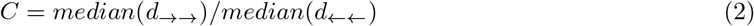

that was the ratio of two medians: the median of the distances between two contigous motifs in the same forward orientation (noted “→ →”) and the median of the distances between two contigous motifs in same reverse orientation (noted “← ←”). Following the 1D genome point of view, the control ratio was supposed to show no difference between the two orientations. Deviations of *C* from 1 might reflect biases in the genome that were not related to CTCF looping.

To assess the significance of ratio *R* (and *C*), we used the Wilcoxon rank-sum test. This test could assess differences of distances even if the distances did not follow a normal distribution.

### 2.2 Validation of *R* as a measure of CTCF-mediated looping

We first studied the ratio *R* using the human genome. For this purpose, the human genome hg38 assembly was used and vertebrate CTCF motifs (JASPAR MA0139.1) were called along the genome. The distance between any two consecutive motifs was computed. To only keep motifs with a higher chance of binding, motifs whose binding scores were lower than a specific quantile threshold were removed. We found that the ratio *R* strongly increased with the binding score and was maximal for a quantile threshold of 80% (Figure 2A). However, the confidence interval of *R* was higher for 80% than for lower quantiles, because too many binding sites were discarded. Thus, as a trade-off, a quantile of 70% was then considered as a threshold for further analyses, because it better allowed comparison of *R* between species with sufficient statistical power (statistical power depends on the number of binding sites).

**Figure 2.**
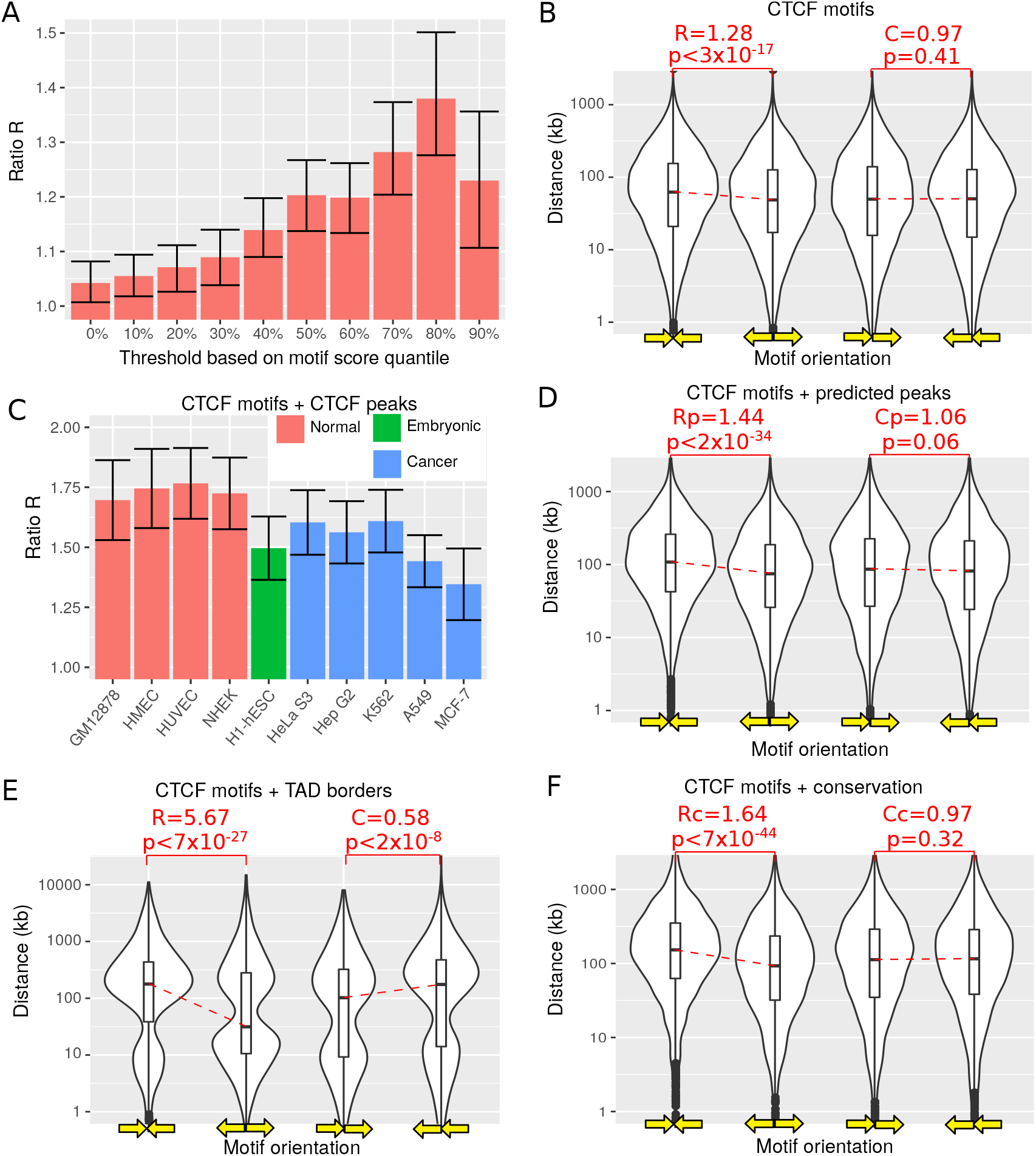
Ratios *R* and *C* computed from the human genome assembly. A) Ratio *R* for different binding score thresholds. B) Distance between consecutive CTCF motifs depending on motif orientation. C) Ratio *R* when accounting for CTCF ChIP-seq data for different cell lines. D) Distance between consecutive CTCF motifs depending on motif orientation, when accounting for predicted CTCF ChIP-seq data. E) Distance between consecutive CTCF motifs depending on motif orientation, when accounting for TAD borders. F) Distance between consecutive CTCF motifs depending on motif orientation, when accounting for conservation score.

We found that the distance between two contigous motifs in convergent orientation was significantly higher than between two contigous motifs in divergent orientation, as expected by the 1D genome point of view of CTCF-mediated looping (*R* = 1.28, Wilcoxon test *p* < 3 × 10^−17^; Figure 2B). In comparison, the distance between two motifs in forward orientation was not significantly different from the distance between two motifs in reverse orientation, as expected by the 1D genome point of view (*R* = 0.97, *p* = 0.41). The bootstrapped distributions of the distance medians were also computed for convergent and divergent motifs, respectively (Supp Fig 1). The two distributions were far apart, reflecting the significant differences of medians. Because the accuracy of the distance between motifs depended on the genome assembly, the ratio was assessed for old and more recent assemblies. As expected, *R* increased with recent assemblies (Supp Fig 2). However, these improvements were very modest, revealing that the assembly version did not have a big impact on the estimation of *R* in human.

We then used CTCF GM12878 ChIP-seq data to remove motifs not bound by CTCF in vivo. The ratio *R* was much higher than previously and very significant (*R* = 1.69, *p* < 5 × 10^−51^; Figure 2C), reflecting the important difference in distance between motifs overlapping CTCF peaks depending on orientation. In vivo information thus helped us to remove false positive motif occurrences and to estimate *R* with more power. We next assessed *R* using CTCF peaks from all ENCODE cell lines. Interestingly, we found that *R* varied depending on cell type. Moreover, *R* was especially low for embryonic stem cells and cancer cells, reflecting lower CTCF looping and thus lower organization of the genome in 3D domains in these cells. However, in practice, only genome assemblies are available for most species and no ChIP-seq data are available. Hence, to circumvent this issue, CTCF ChIP-seq peaks surrounding the motifs were predicted using convolutional neural network learned from human data [1]. This ratio estimated using predicted peaks was noted *R_p_*. The ratio *R_p_* was higher than the one computed from motifs only (*R_p_* = 1.44, *p* < 2× 10^−34^; Figure 2D), revealing the better ratio estimation using peak prediction.

We next filtered motifs located inside 3D domain borders, since those motifs were more likely to influence the 3D genome. For this purpose, we used Arrowhead domains from GM12878 Hi-C data [19]. We extended domain borders to 20 kb on each side and only kept motifs belonging to borders. Accounting for 3D domain borders strikinly improved *R* (*R* = 5.67, *p* < 7 × 10^−27^; Figure 2E). We also filtered motifs located at loop anchors [19]. Again, we extended loop anchors to 20 kb on each side and kept motifs belonging to anchors. Surprisingly, we found a much lower *R* than for 3D domain borders (*R* = 1.77, *p* < 2 × 10^−15^; Supp Fig3).

CTCF binding sites located at 3D domain borders were previously shown to be evolutionary conserved [24]. Hence, we sought to improve *R* computation by discarding non-conserved motifs. This ratio estimated using conservation was noted *R_c_*. This approach greatly improved the ratio (*R_c_* = 1.64, *p* < 7 × 10^−44^; Figure 2F). If both conservation and predicted peaks were used, *R_p_* was even higher (*R_c_* = 1.80, *p* < 5 × 10^−52^). Thus, accounting for conservation score allowed to further improved ratio estimation.

As a control, we computed *R* for *Drosophila* genomes (*melanogaster* and *yakuba*) and *C. elegans*. In *Drosophila melanogaster*, recent high resolution Hi-C data showed the absence of loops mediated by CTCF motifs in convergent orientation [6]. Accordingly, ratio *R* was computed for *melanogaster* and *yakuba* genomes, and were close to one and not significant (dm6: *R* = 0.93, *p* = 0.15; droYak2: *R* = 1.02, *p* = 0.41; Supp Fig 4). In addition, in *C. elegans*, CTCF has been lost during nematode evolution [10]. In agreement, ratio *R* was also close to one and not significant (cell: *R* = 1.02, *p* = 0.22; Supp Fig 4).

Analysis of the human genome thus validated the 1D genome point of view of CTCF-mediated looping. Such looping can be easily estimated from the genomic sequence alone by computing the *R* ratio of distances depending on motif orientation. Moreover, control results revealed the ability of *R* to be equal to one for genomes that are known not to harbour CTCF-mediated loops.

### 2.3 Ratio *R* depends on 3D compartments and isochores

We then computed *R* depending on the underlying genomic and chromatin regions in the human genome. We first investigated if *R* could differ depending on megabase 3D genome compartments, known as A/B compartments, that were shown to divide the genome into gene rich, active and open chromatin (compartment A) and into gene poor, inactive and close chromatin (compartment B) [15]. We found that *R* was greater in compartment B (*R* = 1.40, *p* < 3 × 10^−8^) than in compartment A (*R* = 1.21, *p* < 3 × 10^−8^; Figure 3A), with a slightly significant difference (*p* = 0.03). Accordingly, chromatin loops were larger in compartment B than in compartment A (fold-change=1.4, *p* < 1 × 10^−20^; Supp Fig 5). At high resolution (25 kb), compartments A/B were further shown to be composed of subcompartments A1, A2 (active) and B1, B2, B3, B4 (inactive) [19]. We found that *R* varied between subcompartments. Subcompartments A1 and A2 presented *R* values close to the *R* computed genome-wide (A1: *R* = 1.20, *p* < 4 × 10^−4^; A2: *R* = 1.30, *p* < 4 × 10^−4^; Figure 3B). Conversely, B subcompartments showed high variability of *R*. B1 and B3 showed *R* values greater than the genome-wide *R* (B1: *R* = 1.45, *p* < 4 × 10^−6^; B3: *R* = 1.64, *p* < 2 × 10^−10^; Figure 3B), while B2 and B4 had *R* values that were lower than the genome-wide *R* (B2: *R* = 1.13, *p* = 0.27; B4: *R* = 0.87, *p* = 0.76; Figure 3B). When comparing A and B subcompartments, we found a significant difference between A1 and B3 (*p* = 0.0011). We next analyzed *R* depending on DNA replication timing. We found an *R* value close to the genome-wide value for early replicating regions (*R* = 1.27, *p* < 2 × 10^−4^; Figure 3C), but a high *R* value for late S replicating regions (*R* = 1.57, *p* < 2 × 10^−5^; Figure 3C).

**Figure 3.**
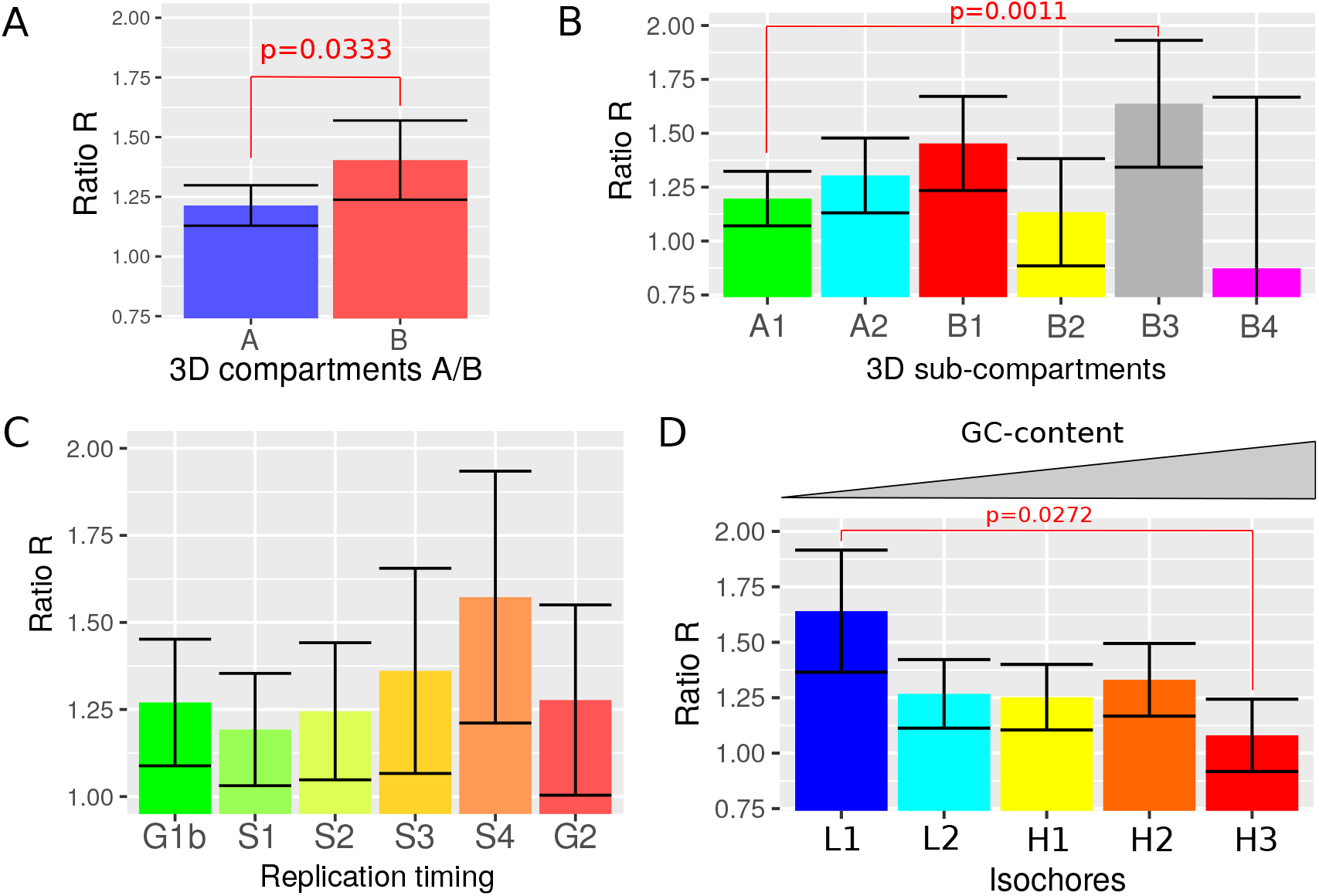
Ratio *R* computed for different chromatin regions in human. A) Ratio *R* estimated for 3D genome compartments A/B. B) Ratio *R* depending on 3D genome subcompartments. C) Ratio *R* and replication timing. D) Ratio *R* depending on GC-content isochores.

Another important feature of the genome is the GC-content that varies considerably along the chromosomes. In particular, the genome was shown to be composed of isochores which are large DNA segments of homogenous GC-content and that were recently shown to be correlated with subcompartments [2,12]. We then computed *R* depending on isochore class (L1, L2, H1, H2 and H3) and observed differences between classes. In particular, L1 isochores (lowest GC-content) showed the highest *R* value (*R* = 1.64, *p* < 5 × 10^−7^; Figure 3D), which was considerably larger than the one estimated genome-wide. Interestingly, L1 *R* value was very close to subcompartment B3 *R* value. Conversely, H3 isochores (highest GC-content) showed the lowest *R* value (*R* = 1.08, *p* = 0.15; Figure 3D), which was lower than the genome-wide *R*.

The *R* ratio thus varied with the underlying genomic and chromatin context, which suggested potential structural and functional roles. Most notably, we found that the ratio *R* is higher in compartment B, in mid-late replication timing regions and in low GC-content isochores, which were associated with heterochromatin.

### 2.4 CTCF looping in mammals

We then estimated *R* for available mammal genomes. Because the accuracy of *R* estimation depended on the number of motif pairs, we computed *R* for genomes with a sufficient number of pairs (> 8000). We found that all mammals presented an *R* value that was superior to one and significant (Figure 4A). The tasmanian devil and the pika presented the highest values (*R* > 1.5), whereas the horse and the guinea pig showed the lowest values (*R* close to 1.2). It was very interesting to see that *R* estimation could be significantly different from zero even for assemblies whose qualities were much lower than hg38, such as papAnu2 (scaffold *N*_50_ = 586 kb, scaffold *L*_50_ = 1481; *R* = 1.37, *p* < 9 × 10^−22^) and ornAna2 (scaffold *N*_50_ = 959 kb, scaffold *L*_50_ = 309; *R* = 1.44, *p* < 7 × 10^−18^).

**Figure 4.**
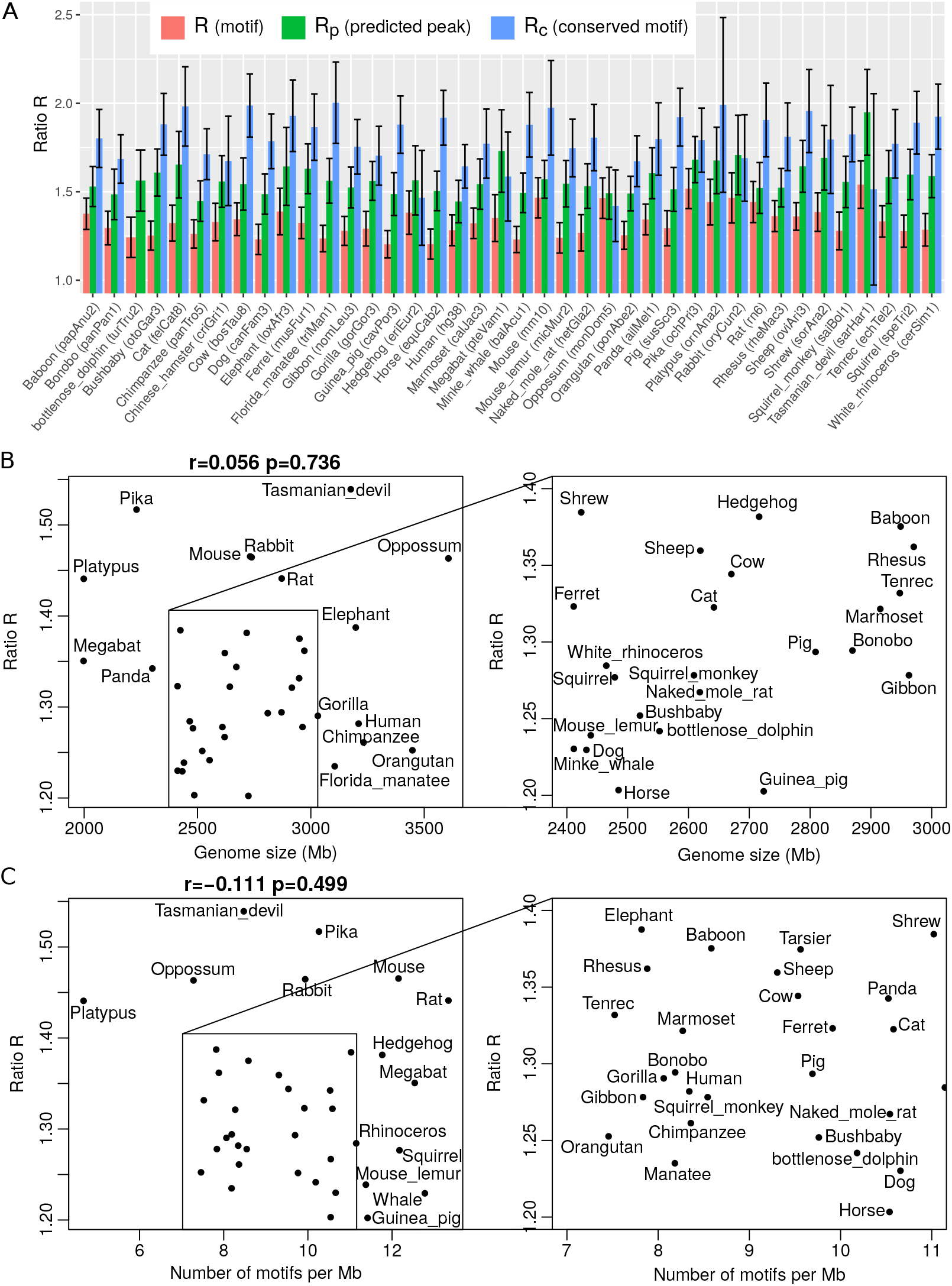
Ratio *R* computed from mammal genomes. A) Ratios *R*, *R_p_* and *R_c_* computed from all mammal genome assemblies. B) Ratio *R* versus genome size. C) Ratio *R* versus motif density (number of motifs per Mb).

We also predicted CTCF ChIP-seq peaks surrounding the motifs, and estimated *R_p_*. The ratio *R_p_* was superior to *R* estimated from motifs only (Figure 4A). Interestingly, although the convolutional neural network we used was trained from human data, it could dramatically increase the ratio for most species. For instance, *R_p_* was higher than *R* for the dog (canFam3: *R* = 1.20, *R_p_* = 1.49, 24% increase) and even for the platypus (ornAna2: *R* = 1.44, *R_p_* = 1.68, 17% increase). We also filtered conserved motifs and computed *R_c_*. The ratio *R*_c_ was even higher than *R_p_* for most species. For example, *R_c_* were higher than *R* and *R_p_* for the dog (canFam3: *R* = 1.20, *R_c_* = 1.79, 49% increase) and the platypus (ornAna2: *R* = 1.44, *R_p_* = 1.99, 38% increase). However, a major drawback of *R_c_* and *R_p_* was their larger confidence intervals, and that is the reason why we kept *R* for further analyses.

We next investigated if *R* was influenced by the genome size, which could explain the observed differences of *R* between species. No significant correlation was found between the genome size and *R* (Figure 4B), confirming that *R* was not biased by the genome size, and thus allowing *R* comparison between species. We also assessed if *R* was influenced by the density of motifs in the genome (number of motifs per Mb), and no significant correlation was found (Figure 4C). For instance, the platypus and rat genomes presented an *R* value around 1.45, but contained 4.66 motifs per Mb and 13.33 motifs per Mb, respectively.

The difference of *R* between two species could then be tested using a permutation test. For instance, the Tasmanian devil and human genomes which have a similar size were compared (both around 3.2 Gb). The tasmanian devil genome (*R* = 1.54) had a significantly higher *R* than the human (*R* = 1.28) (*p* = 0.0004), which could reflect a stronger CTCF looping in the Tasmanian devil genome. Using predicted peaks, the tasmanian devil genome (*R* = 1.95) also had a significantly higher *R* than the human genome (*R* = 1.44) (*p* < 1 × 10^−5^). Because this difference might be due to potential assembly inaccuracy of the Tasmanian devil genome assembly, we then compared two accurate assemblies: the human genome assembly hg38 and the mouse genome assembly mm10. We found that the mouse genome (*R* = 1.47) had a significantly higher *R* than the human genome (*R* = 1.28) (*p* = 0.0028).

The *R* ratio can thus be used to study the 3D genome organization in CTCF loops in mammals even for species whose Hi-C data were not available. Moreover, we found important differences of *R* between mammals, some of them were statistically significant. For instance, we found that species that were evolutionary distant, such as the human and the Tasmanian devil, presented an important difference of *R*.

### 2.5 Phylogenetic analysis of CTCF looping in vertebrates

We then estimated *R* for vertebrate species in order to investigate differences between mammals, reptiles, amphibians, and fishes. Ratios *R* were plotted on the phylogenetic tree to investigate the potential link between CTCF looping and evolution (Figure 5A). Among the vertebrates, the jaw fishes presented very high *R* values, especially the tetraodon (tetNig2: *R* = 1.65, *p* < 2 × 10^−27^) and fugu (fr3: *R* = 1.59, *p* < 4 × 10^−60^). Surprisingly, the zebrafish presented a low *R* = 0.98, which was inconsistent with recent Hi-C results supporting loop formation by CTCF in convergent orientation [14]. In addition, the amphibian xenopus showed a very high *R* value (xenTro7: *R* = 1.63, *p* < 3 × 10^−79^). Interestingly, using peak prediction models trained on human data, the Rp values were even higher: tetraodon (Rp = 1.84, *p* < 8 × 10^−90^) and *Xenopus* (*R_p_* = 1.85, *p* < 2 × 10^−24^). Lampreys which are jawless fishes that diverged from the jawed vertebrate lineage more than 500 million years ago also revealed a significant ratio (*R* = 1.30, *p* < 7 × 10^−9^), supporting the ancient establishment of CTCF looping prior to vertebrates [9].

**Figure 5.**
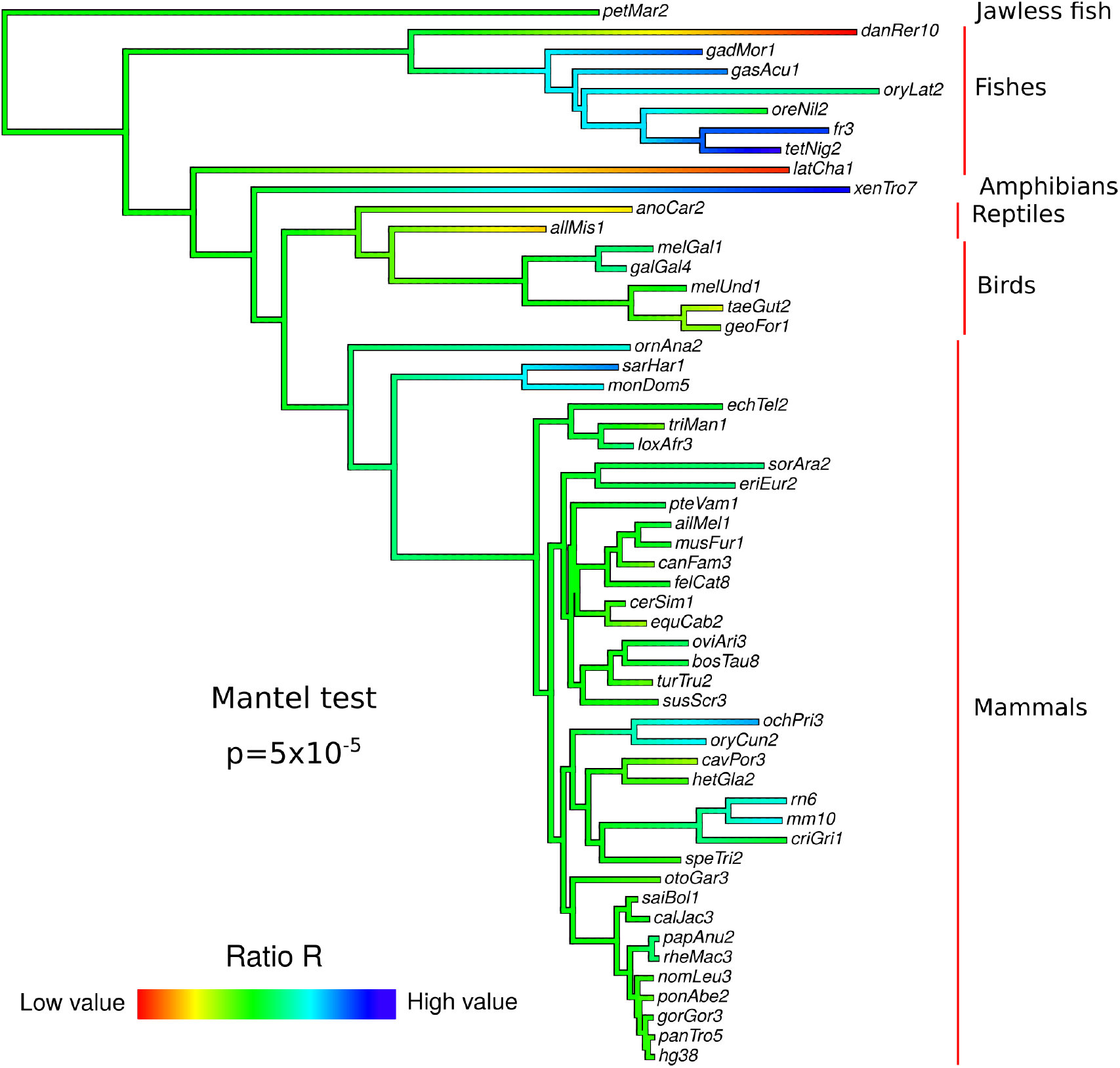
Phylogenetic analysis of *R* in vertebrates. Ancestral *R* reconstruction was done using maximum likelihood inference.

The different assemblies did not have the same quality, which thus introduced some inaccuracy in the estimation of *R*, especially for species that were recently sequenced (those with a assembly number close to one). Despite *R* inaccuracy due to heterogenous assembly quality, we found that evolutionary close species tended to have a similar *R* value (Mantel test *p* = 5 × 10^−5^), revealing conservation of *R* among species (Figure 5A). For instance, two relatively close species in the tree, the rat (rn6: *R_p_* = 1.44, *p* < 1 × 10^−27^) and the mouse (mm10: *R_p_* = 1.47, *p* < 3 × 10^−36^) presented very similar *R* values (*p* = 0.61). Hence, ancestral *R* reconstruction could be carried out (Figure 5A). It revealed that a large *R* value was acquired in the common ancestor of the rat and the mouse (Supp Fig 6). Similar findings were observed for the American pika (ochPri3) and the European rabbit (oryCun2), and also for the Tasmanian devil (sarHar1) and the opossum (monDom5).

Another important parameter contributing to CTCF looping is the CTCF motif density in bilaterian genomes [9]. Hence, we estimated CTCF motif density in vertebrates and observed a strong conservation (Mantel test *p* < 1 × 10^−5^; Supp Fig 7). Jaw fishes showed high motif densities, such as the fugu (fr3: 40.58 motifs/Mb). Conversely, birds showed very low motif densities, such as the chicken (galGal4: 7.05 motifs/Mb). Mammals presented varying densities, for instance 4.66 for the platypus (ornAna2) and 21.4 for the Chinese hamster (criGri1). Among mammals, we observed very homogenous clades, such as primates, whose *R* varied from 7.45 to 9.76. Moreover, we found that CTCF motif density was evolutionary conserved (Mantel test *p* < 1 × 10^−5^), which suggested that ancestral motif density reconstruction could be done. Inference of ancestral density uncovered interesting results, such as the low density for the primate ancestor as compared to the higher density for the muridae ancestor.

Results revealed the evolutionary conservation of *R* among vertebrates. *R* thus represented a useful tool to study 3D genome evolution, in addition to CTCF motif density. The two parameters could be used to study CTCF looping in ancestral genomes by using ancestral character reconstruction.

## 3 Conclusion

In this paper, we propose a novel approach to study the 3D genome evolution in vertebrates using the genomic sequence only, without the need of costly and challenging Hi-C data to produce. Therefore, the approach allows a comprehensive analysis of vertebrates whose genome assemblies are now available and whose number will exponentially increase with large sequencing projects such as the Vertebrate Genomes Project (VGP) aiming to sequence 66,000 extant vertebrate species. The proposed approach is very simple and makes very few assumptions. It relies on the CTCF motif which is known to be conserved across vertebrates and the CTCF looping model that implies a 1D genome point of view where convergent motifs are expected to be more distant than divergent motifs. The approach can be further improved by using predicted CTCF ChIP-seq peaks or by using the conservation score surrounding the CTCF motif, reflecting strong conservation of the DNA context surrounding CTCF motifs in vertebrates, especially for mammals. Using the human genome as a reference, we validate the 1D genome point of view and demonstrate that the ratio of distances between convergent and divergent motif pairs (ratio *R*) can quantify CTCF looping. These results reflect strong evolutionary constraints encoded in the genome that are associated with the 3D genome organization.

The proposed approach also uncovers a number of results. We found that *R* varies with the underlying genomic and chromatin regions, such as 3D compartments and sub-compartments, isochores and replication timing, which supports a potential structural and functional role of *R*. In particular, *R* encodes higher CTCF looping in heterochromatin regions. Moreover, the analysis of *R* combined with CTCF ChIP-seq peaks showed a lower value for *R* in cancer and embryonic cells compared to normal cell lines. Thus, depending on the cell state, *R* can be modulated by CTCF binding in vivo, thereby regulating CTCF looping strength. Regarding *R* in different species, we show most notably that *R* is evolutionary conserved among vertebrates. Species that are phylogenetically close tend to have a ratio that is closer than species that are phylogenetically far. Among vertebrates, several fishes and amphibians show the highest ratio, whereas reptiles show low values. In mammals, ancestral character reconstruction reveals that the genome of the ancestor of the rat and mouse likely evolved to have a high *R* value. A previous study showed the linear divergence of CTCF binding sites with evolutionary distance, and the birth of new genes associated with the birth of new CTCF binding sites [16]. Here, our approach reveals that the distance between convergent motifs which underlies CTCF looping and TAD organization evolves over time between vertebrates, and thus represents an important factor contributing to 3D genome evolution.

There are several limitations of the proposed approach. First, the estimation of distances between CTCF motifs depends on the genome assembly quality. Thus, for draft genomes, it is likely that the *R* ratio will not be accurately estimated, especially when scaffolds are small. Second, deep learning models can be used to improve *R* for species without any available ChIP-seq data, but the models were learned from human data and thus CTCF peak prediction is expected to be less accurate for species that are very distant from human, leading to *R* underestimation. Third, we found a non-significant *R* value for the zebrafish (danRer10) which is in contradiction with recent Hi-C data [14], thus revealing the inadequacy of *R* for certain vertebrate species.

## 4 Materials and Methods

### 4.1 Hi-C data, compartments, subcompartments and TADs

In human, we computed compartments A/B using Juicer Tools [5]. For this purpose, we used publicly available Hi-C data from GM12878 cells from Gene Expression Omnibus (GEO) accession GSE63525 [19]. For subcompartments, we downloaded the genomic coordinates from GEO GSE63525. For TAD borders and loop anchors, we downloaded respectively Arrowhead domains and HiCCUPS loops called from GM12878 Hi-C data from GEO GSE63525.

### 4.2 Isochores

In human, we called isochores using isoSegmenter program on hg38 assembly [3].

### 4.3 Replication timing

In human, we used GM12878 Repli-seq from ENCODE [22].

### 4.4 CTCF motif calling

We used the vertebrate CTCF motif position frequency matrix MA0139.1 from the JASPAR database (http://jaspar.genereg.net/). We scanned CTCF binding sites on the following genome assemblies: ailMel1, allMis1, anoCar2, apiMel2, aplCal1, aptMan1, balAcu1, bosTau8, braFlo1, calJac3, calMil1, canFam3, cavPor3, cell, cerSim1, choHof1, criGri1, danRer10, dipOrd1, dm6, droYak2, echTel2, equCab2, eriEur2, felCat8, fr3, gadMor1, galGal4, gasAcu1, geoFor1, gorGor3, hetGla2, hg38, latCha1, loxAfr3, macEug2, melGal1, melUnd1, micMur2, mm10, monDom5, musFur1, myoLuc2, nomLeu3, ochPri3, oreNil2, ornAna2, oryCun2, oryLat2, otoGar3, oviAri3, panPan1, panTro5, papAnu2, petMar2, ponAbe2, proCap1, pteVam1, rheMac3, rn6, saiBol1, sarHar1, sorAra2, speTri2, strPur2, susScr3, taeGut2, tarSyr2, tetNig2, triMan1, tupBel1, turTru2, vicPac2, xenTro7. For this purpose, we used MEME FIMO program with default parameters (http://meme-suite.org/doc/fimo.html).

### 4.5 CTCF ChIP-seq peak

In human, we used CTCF ChIP-seq peaks for several cell lines from ENCODE (https://genome.ucsc.edu/encode/).

### 4.6 Deepbind

To improve binding predictions for CTCF, we used deepbind to predict binding on the 500 base region surrounding motif occurrence (http://tools.genes.toronto.edu/deepbind/). We used the deepbind model trained on CTCF ChIP-seq data, noted D00328.018.

### 4.7 Conservation score

We computed the average conservation score of the 50 bases surrounding the CTCF binding sites using hg38 phastCons scores from UCSC Genome Browser (https://genome.ucsc.edu/). For other assemblies, we liftovered hg38 phastCons scores.

### 4.8 Permutation test to compare two *R* values

A permutation test was devised to test the difference of *R* between two species 1 and 2. For each permutation, the test consisted in shuffling species labels between motif pairs, and then in computing *d_R_* = |*R*_1_ – *R*_2_|. The p-value was computed as the percent of *d_R_* values from permutations that were superior or equal to the observed *d_R_* value.

### 4.9 Code availability

The functions to compute *R*, to test difference of *R*, and to reconstruct the ancestral *R* were developed in the R language. There are available at https://github.com/morphos30/PhyloCTCFLooping under Apache License 2.0.

## 5 Supplementary Figures

**Supp Fig 1:**
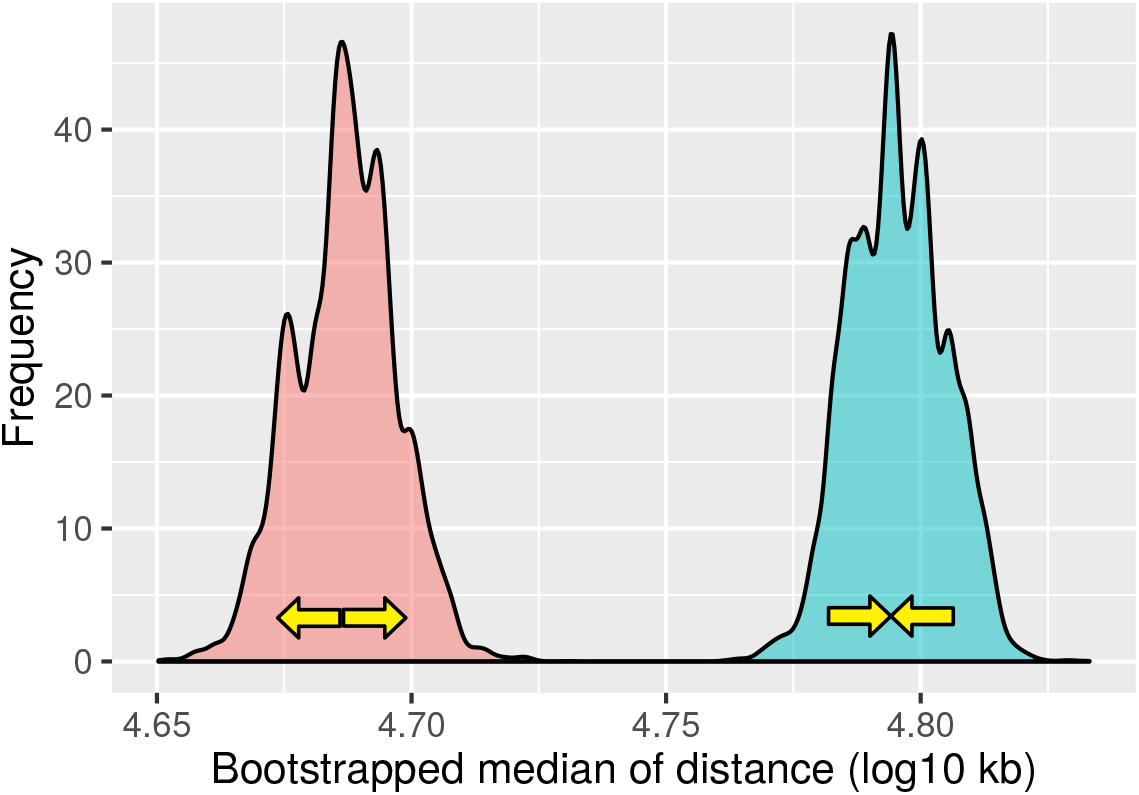
Distribution of the bootstrapped median of distance (log10 kb).

**Supp Fig 2:**
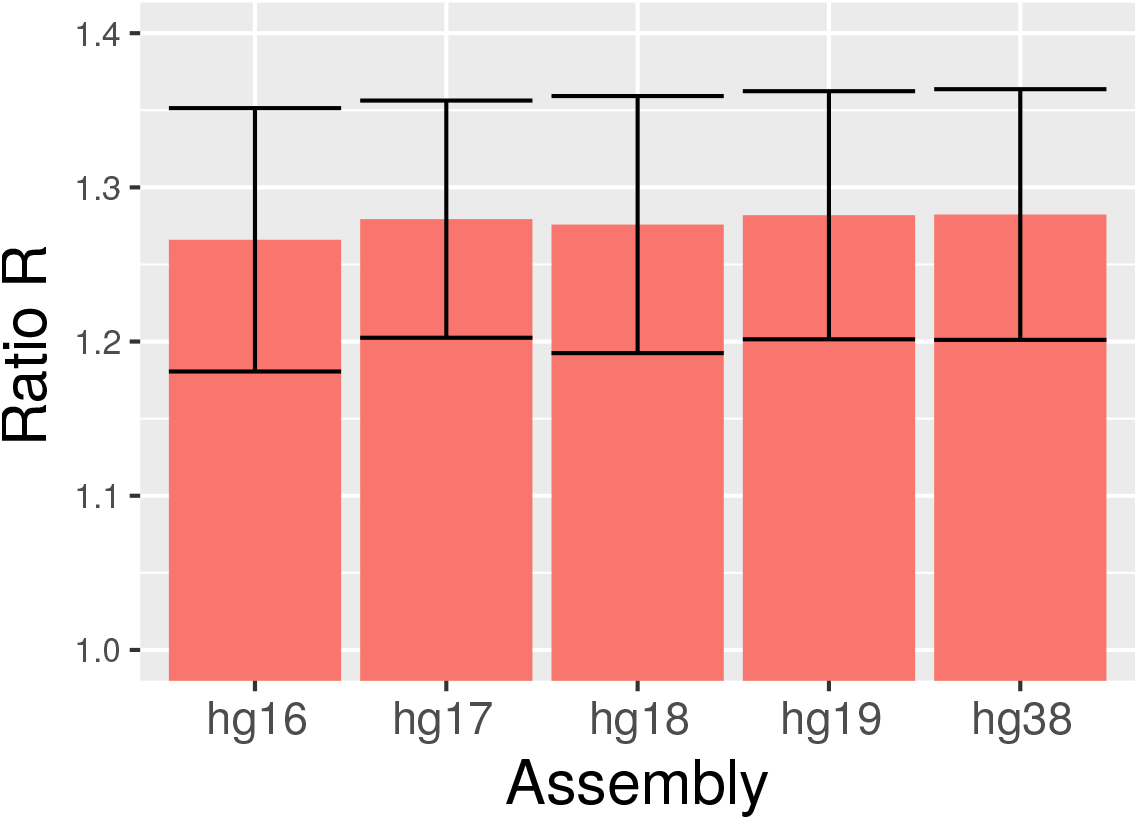
Ratio *R* computed from different human genome assemblies.

**Supp Fig 3:**
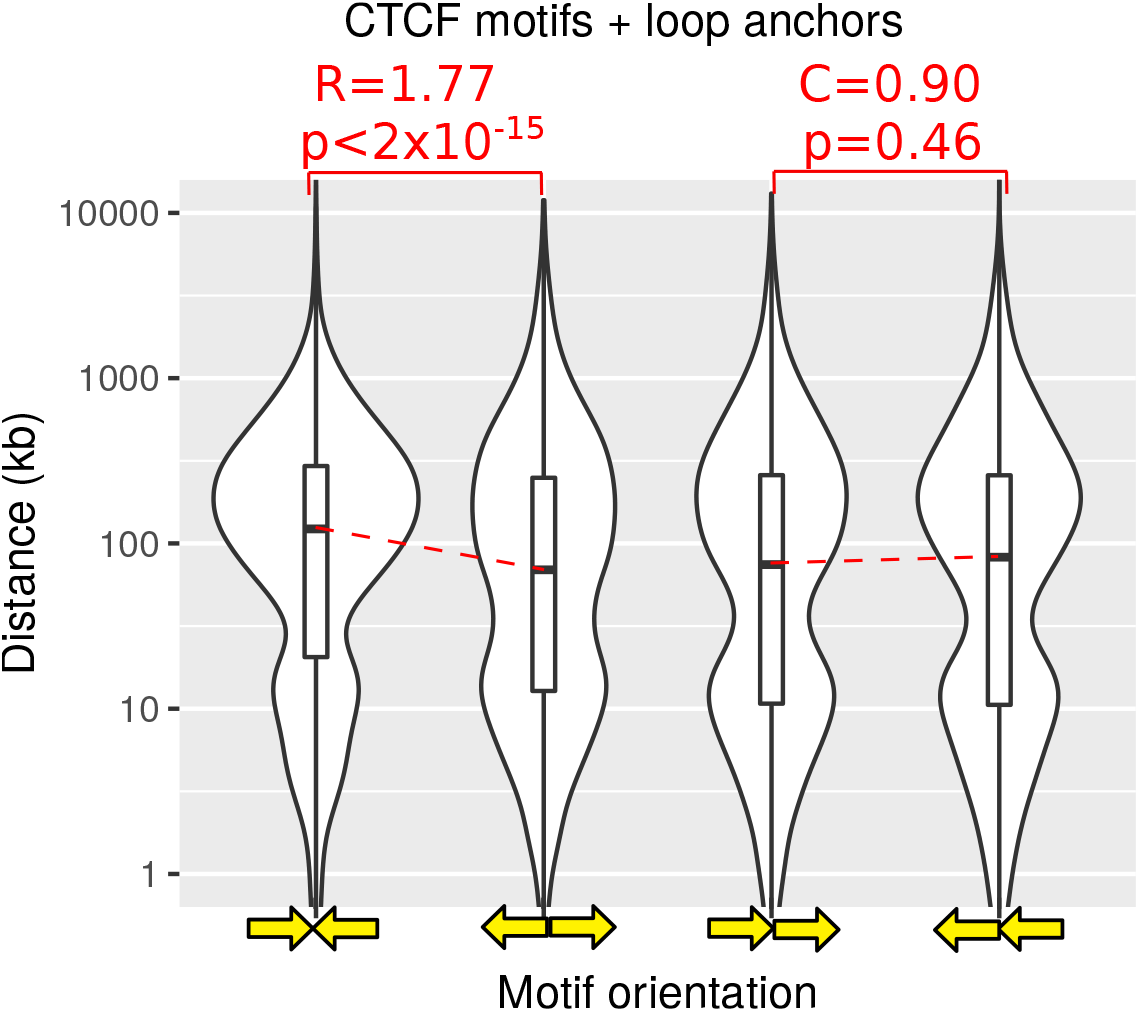
Ratios *R* and *C* computed from the human genome assembly, when accounting for Hi-C loop anchors.

**Supp Fig 4:**
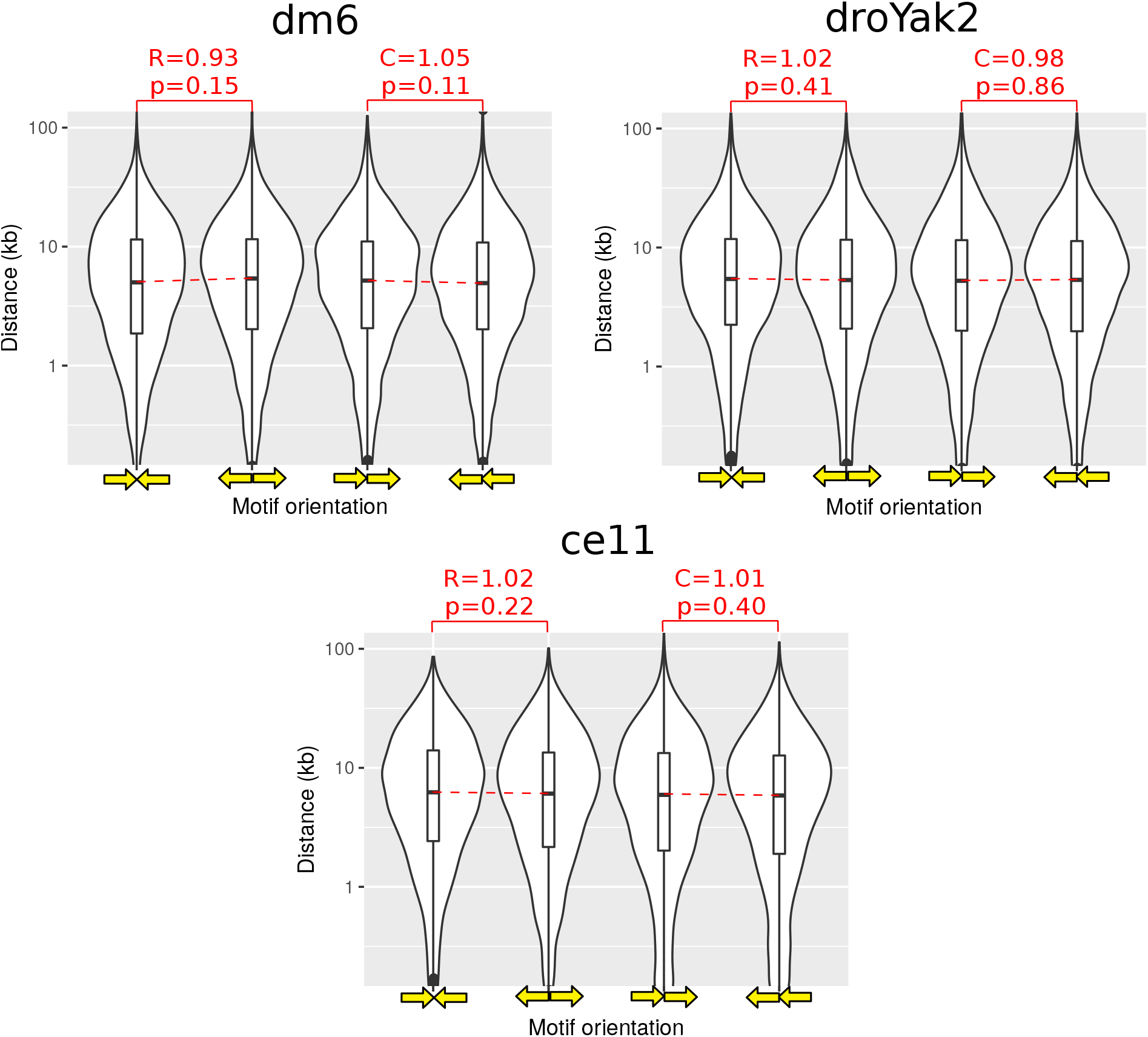
Ratios *R* and *C* computed from *Drosophila melanogaster* dm6 and *yakuba* droYak2 genome assemblies, and from *C. elegans* cell genome assembly.

**Supp Fig 5:**
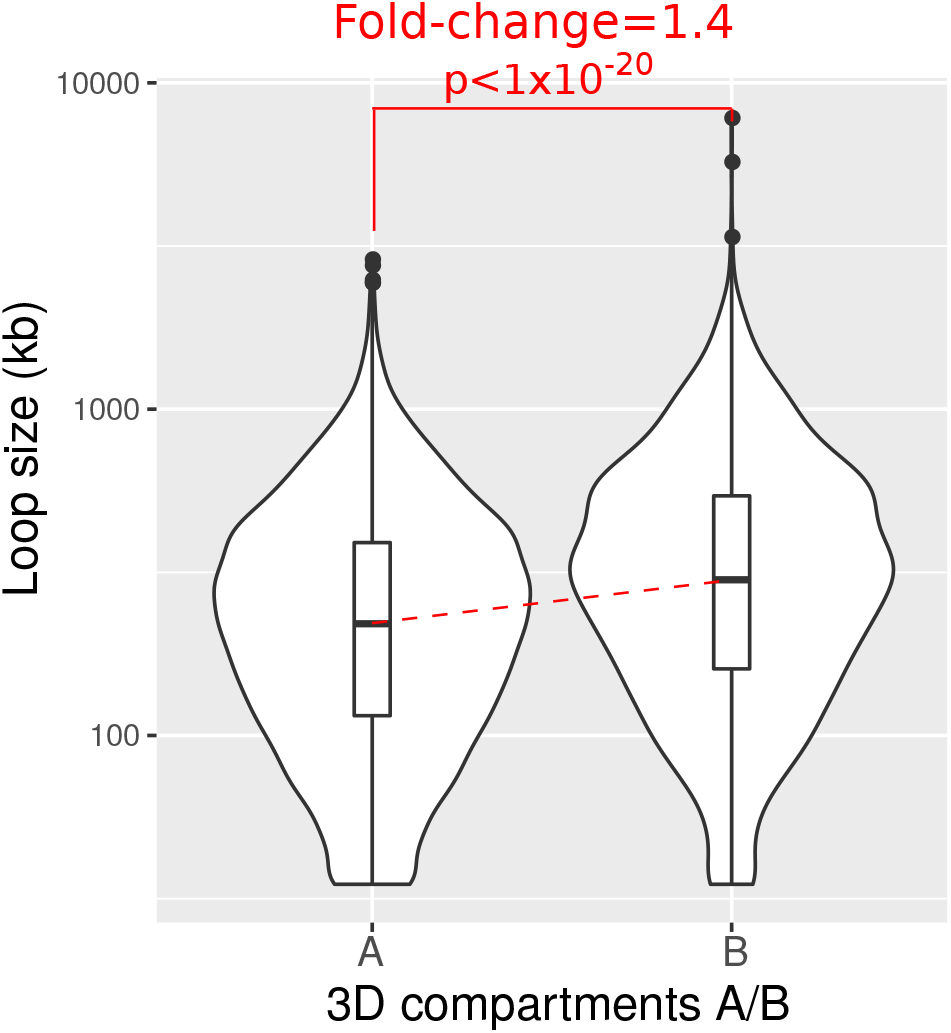
Chromatin loop size depends on 3D compartments A and B in human.

**Supp Fig 6:**
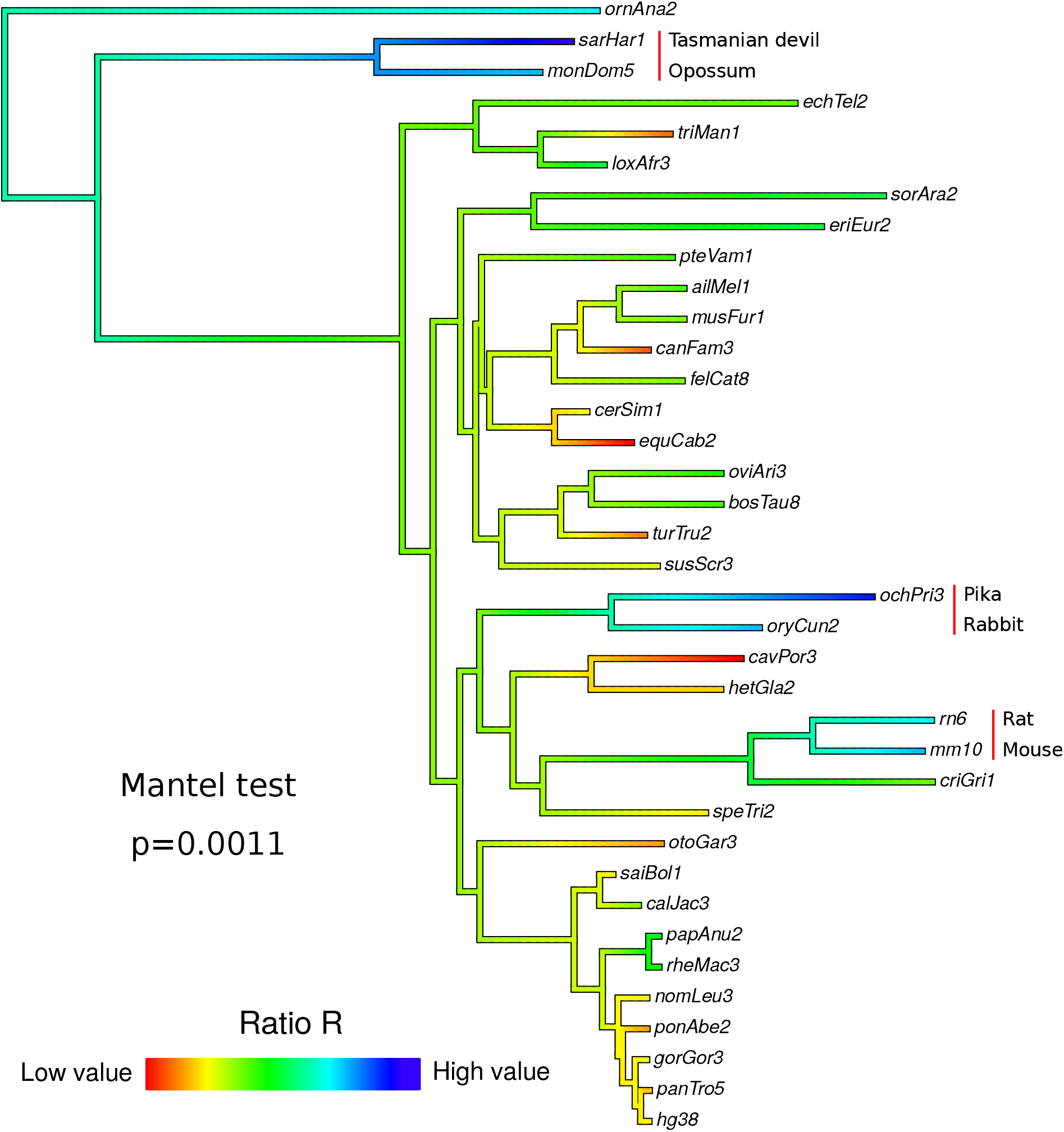
Phylogenetic analysis of ratio *R* in mammals. Ancestral ratio *R* reconstruction was done using maximum likelihood inference.

**Supp Fig 7:**
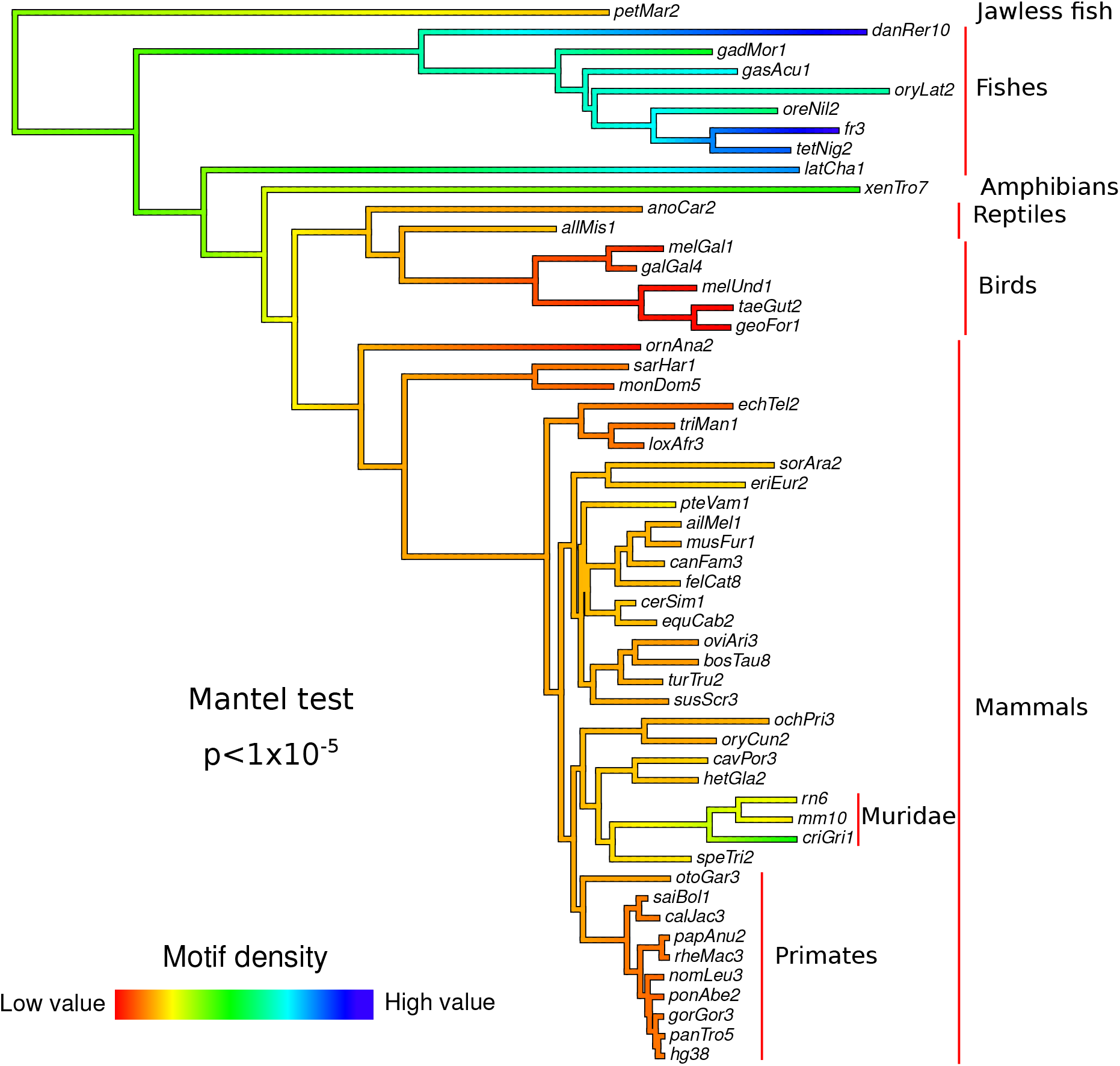
Phylogenetic analysis of CTCF motif density in vertebrates. Ancestral motif density reconstruction was done using maximum likelihood inference.

